# mb-PHENIX: Diffusion and Supervised Uniform Manifold Approximation for denoising microbiota data

**DOI:** 10.1101/2022.06.23.497285

**Authors:** Padron-Manrique Cristian, Vázquez-Jiménez Aarón, Esquivel-Hernandez Diego Armando, Martinez Lopez Yoscelina Estrella, Neri-Rosario Daniel, Sánchez-Castañeda Jean Paul, Giron-Villalobos David, Resendis-Antonio Osbaldo

## Abstract

**Motivation:** Microbiota data suffers from technical noise (reflected as excess of zeros in the count matrix) and the curse of dimensionality. This complicates downstream data analysis and compromises the scientific discovery’s reliability. Data sparsity makes it difficult to obtain a well-cluster structure and distorts the abundance distributions. Currently, there is a rised need to develop new algorithms with improved capacities to reduce noise and recover missing information.

**Results:** We present mb-PHENIX, an open-source algorithm developed in Python, that recovers taxa abundances from the noisy and sparse microbiota data. Our method deals with sparsity in the count matrix (in 16S microbiota and shotgun studies) by applying imputation via diffusion onto the supervised *Uniform Manifold Approximation Projection* (sUMAP) space. Our hybrid machine learning approach allows the user to denoise microbiota data. Thus, the differential abundance of microbes is more accurate among study groups, where abundance analysis fails.

**Availability:** The mb-PHENIX algorithm is available at https://github.com/resendislab/mb-PHENIX. An easy-to-use implementation is available on Google Colab (see GitHub)

**Supplementary information:** Supplementary data are available at *Bioinformatics* online.

## 1 Introduction

Advances in high-throughput technologies have been essential for exploring the association between microbiota and human diseases. However, analyzing microbiota data is a challenge due to many sources of technical noise (i.e., DNA extraction, PCR, and library preparation, etc.,). Consequently, the taxa count matrix contains many zeros which drives a zero-inflated abundance distribution (*Jiang et al., 2015*). This noisy effect is a ubiquitous phenomenon in the sequencing methods 16S rRNA and whole metagenome shotgun (W-MS). Another complication in microbiome data analysis is their high dimensionality. Differences in a high-dimensional space are homogeneous leading to associate samples as similar in an inaccurately manner. To deal with the high-dimensionality and obtain feature insights that explain the heterogeneity of a biological system, different unsupervised dimensionality reduction methods approaches have been proposed such as PCA, MDS and t-SNE (*Armstrong et al., 2021*). Recently, UMAP (a superior manifold learning method) has been used to reveal complex patterns and resolve visualization artifacts in microbiota data (*Armstrong et al., 2021*). Therefore, supervised UMAP (sUMAP) for high-heterogeneous data has been suggested (*McInnes et al., 2018*), as the microbiota data. Here, we present mb-PHENIX, a data imputation algorithm via diffusion on supervised UMAP. This being a novel approach to recover missing data for microbiota data. This method significantly improves the imputation of microbiome data compared to other methods recovering relevant information unmasking the biological processes (see, supplementary Text S1).

## 2 Workflow of the supervised mb-PHENIX Algorithm

The input of mb-PHENIX is an abundance matrix where the columns are the taxa (or ASV/OTUs ID) and rows are samples (Fig S3). Data can be or not be pre-processed and normalized using standard workflows, (see, supplementary Text S2). However, we acknowledge that normalization methods for microbiota data is a separate project. For the sUMAP embedding, we required the categorical information of the samples in study, for example: cases and controls. The main idea of supervised embedding is to map data in different classes in a low dimensional space as far apart as possible, maintaining the internal class structure and the inter-classes relationships (see supplementary Text S1 and S3 and the Fig S4). Therefore, we computed a distance matrix from sUMAP space. Then, we obtained a stochastic Markov transition matrix (*M*) representing a sample-to-sample transition probability. After that, the diffusion process occurs. *M* is exponentiated to a discrete ***t*** value. This parameter simulates the diffusion process of the random walk along the reduced multidimensional sUMAP space. Finally, the missing taxon data is recovered by the matrix multiplication between *M*^*t*^ and the sparse taxa matrix used as input. As a result, the missing data are estimated in terms of those data points with the nearest neighborhoods. In summary, mb-PHENIX recovers data based on the nearest neighbors in the manifold obtained by the diffusion process on the sUMAP space, see general workflow in Fig S2. Just to point out, mb-PHENIX is a supervised variant from unsupervised sc-PHENIX method (*Padron-Manrique et al*., *2022*). The difference lies that sc-PHENIX uses UMAP in an unsupervised manner and mb-PHENIX does not.

## 3 Applications

The With the purpose to exemplify our method, we applied mb-PHENIX onto 16S rRNA microbiota data from a cohort with individuals with different degrees of type 2 diabetes(*Diener et al., 2019*). In Figure 1, we depict the abundance matrix of the highly variable taxa in three conditions: without imputation (A), imputed with unsupervised mb-PHENIX(B), and supervised mb-PHENIX(C). Also, we imputed data with mbImpute, a specialized imputation method for microbiome data (*Jiang et al., 2015*) and MAGIC (*van Dijk et al., 2018*), an imputation method for scRNA-seq. However, these methods do not not recover cluster structure of the data (Fig S7 to S9).

**Fig. 1.**
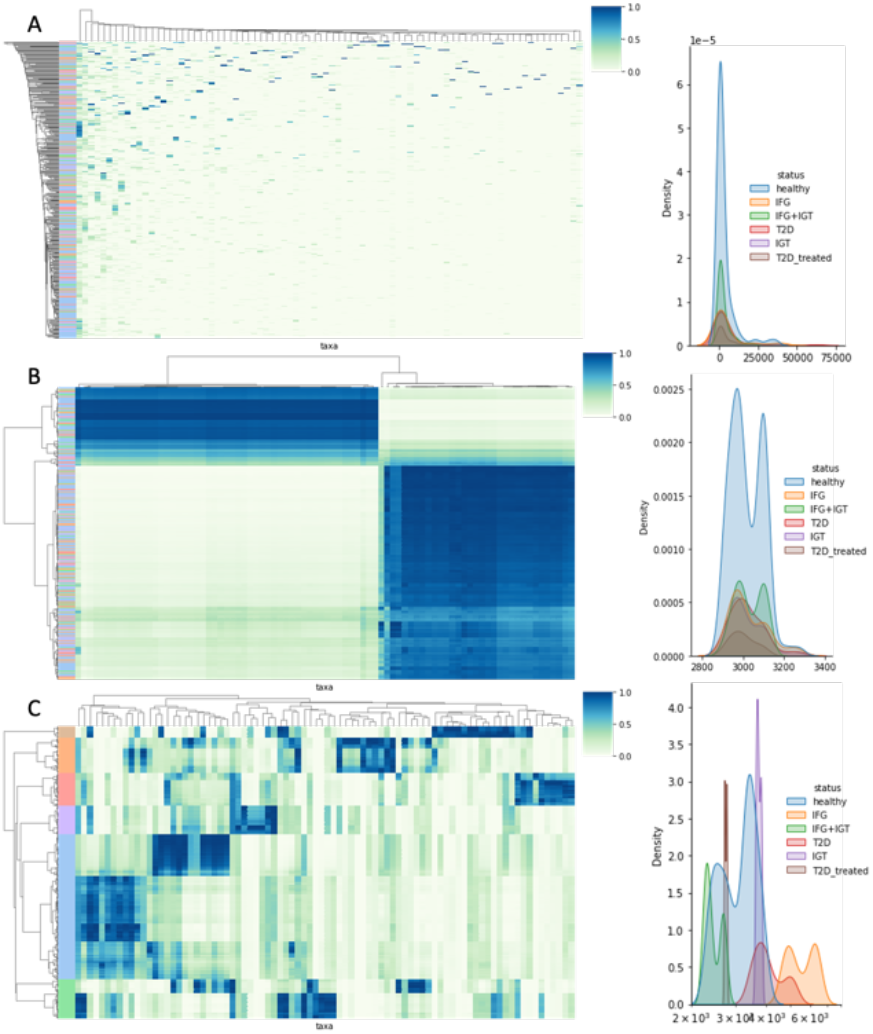
The method mb-PHENIX reveals multinomial distribution of taxa in the pro-gregression of T2D microbiota data. Left: plots of the raw and imputed taxa matrix as a hierarchically-clustered heatmap. We visualize the most variable taxon (95 percentile) of non-imputed data (A) unsupervised imputated (B) and supervised imputated(C) data with mb-PHENIX. Right: Histograms by state clusters for *Akkermanssia* taxon calculated using Kernel Density Estimation (KDE) on the data before (A) and after unsupervised (B) and supervised (C) mb-PHENIX imputation. After supervised mb-PHENIX, we observed separated multimodal distributions per taxa, with different transition states of TD2 clusters represented by different peaks, consistent with the known taxon in these microbiome subsets. TD2= Type 2 Diabetes, IFG= Impaired Fasting Glucose, IGT = Impaired Glucose Tolerance, T2D_treated = Treated Diabetes, IFG + IGT = Impaired Fasting Glucose and Impaired Glucose Tolerance.

We observed that the non-imputed count matrix does not recover any cluster structure and taxon has a unimodal distribution centered at 0 (right side of Fig 1A). Similar results are obtained with mbImpute and MAGIC the unimodal distribution is near to 0 (Fig S7 to S10). The unsupervised imputed count matrix generates a bimodal distribution(Fig 1B). Contrastingly, after we applied the supervised mb-PHENIX onto the count matrix (sUMAP space), we observed a multi-modal distribution per taxon (Fig 1C). Thus, the count matrix obtained from mb-PHENIX leads to obtaining the differential abundances among conditions better than other methods.

## 4 Discussion and conclusion

We present a novel imputation method that contributes to recover the lost information in the abundance matrix of microbiota. Although similar results were obtained from *Diener et al 2019*, new differential abundance taxa were unmasked. The presented method is integrated in an open source developed in Python. As we showed, the combination of Supervised UMAP with imputation via diffusion brings an improved approach to recover missing information for microbiota data (Fig 1C, S5 and S6).

## Acknowledgements

OR-A thanks the financial support from CONACYT (Grant Ciencia de Frontera 2019, FORDECYT-PRONACES/425859/2020), PAPIIT-UNAM (IA202720), and an internal grant from the National Institute of Genomic Medicine (INMEGEN, México). PM-C is a doctoral student from Programa de Doctorado en Ciencias Biomédicas, Universidad Nacional Autónoma de México (UNAM) and received fellowship to CVU 855825 from CONACYT, México. This paper is part of the doctoral thesis and the requirements to obtain the degree of Doctor in Science to PM-C. YEM-L is a doctoral student from Programa de Doctorado en Ciencias Médicas, Odontológicas y de la Salud by Universidad Nacional Autónoma de México (UNAM) and received fellowship to CVU 629384 from CONACYT. DAE-H is a postdoctoral research associate at INMEGEN and received CONACyT fellowship to CVU 420693. DN-R is a student from the Master in Sciences program in Ciencias Bioquímicas, UNAM and received CONACyT fellowship to CVU 1083211. JPS-C is a student from the Master in Sciences program in Ciencias Bioquímicas, UNAM and received CONACyT fellowship to CVU 1005702. DG-V is a student from the Master in Sciences program in Ciencias Bioquímicas, UNAM and received CONA-CyT fellowship to CVU 1083058.

## Supplementary Figures and Text

### S1 mb-PHENIX assumption

The supervised based imputation of mb-PHENIX is based on the assumption that data has high amounts of missing data represented as zeros, misleading to find a well-clustered structure on the microbiota data. In addition, microbiota data is highly dimensional and using unsupervised reductional methods such as PCA or any multidimensional scaling variants does not allow finding well-defined patterns with traditional methods (*Fouquier et al*., *2021*). mb-PHENIX is ideal for datasets where samples have the same experimental sampling process and sequencing methodology. However, due to technical noise, available methods fail to find topological patterns and information about the differential abundance. mb-PHENIX solves this situation by imputing similar samples based on the nearest neighbors of the supervised embedding with UMAP (sUMAP). The goal of sUMAP is to map different classes in the low dimensional space as far apart as possible, while at the same time maintaining the internal class structure and the inter-class relationships. Consequently, missing information of abundances is recovered among similar samples mapped on the sUMAP space via diffusion. The diffusion process is the same as reported for sc-PHENIX (*Padron-Manrique et al*., *2022 and van Dijk et al*., *2018*), see Fig S2.

### S2 Preprocessing

The technologies 16S rRNA gene and whole metagenome shotgun (W-MS) sequencing data share the same problems such as data sparsity (due to technical noise) and the curse of dimensionality (due to their high-dimensionality, every taxon is a dimension). Both technologies are processed into the same data structure containing abundances of microbes in microbiome samples: a taxon count matrix with rows as microbiome samples (subjects or individuals) and columns as taxa (i.e., OTUs or ASVs for 16S rRNA data and species for WGS data), and each entry corresponds to the number of reads mapped to a taxon in a microbiome sample. We recommend filtration detected reads that do not appear frequently on samples.

However, to compare taxa counts among samples, normalization methods are required. We recommend doing a library size normalization. However, we acknowledge that normalization methods for microbiota data is a separate project. Hence, we refer users to benchmark papers of normalization (Calgaro et al., 2020 and McKnight et al., 2019) if users consider normalizing the data.

### S3 sUMAP space transformation and parameter selection handling

mb-PHENIX needs two essential inputs: the count matrix (normalized or not) to be imputed and the supervised reductional dimension representing the high-dimensionality of the count matrix data. Basically, mb-PHENIX smooths data based on nearest neighbors on the low-dimensional manifold captured by the exponentiated Markov matrix. The user also can choose the classical use of UMAP in an unsupervised manner (if there is a well-cluster structure on the data). But for high-heterogenous microbiota data, the count matrix preprocess is embedded by supervised UMAP. For the supervised dimension reduction, UMAP takes the labels as a separate metric space (with a categorical distance on it). Then tries to intersect the fuzzy-simplicial set of the high-dimensional data and labels(categorical) data together (McInnes et al., 2018). It is important to make labels assigned as integers and not string data types. For example, ‘T2D’ needs to be 0 and ‘Control’ needs to be 1 for sUMAP fit.

For sUMAP parameters:

For more understanding of how to embed data in a supervised manner with UMAP please refer to the UMAP documentation (https://umaplearn.readthedocs.io/en/latest/supervised.html)

#### target_weight

For supervised embedding with UMAP, the parameter is a weighting factor between data topology and target topology. A value of 0.0 weights predominantly on data, a value of 1.0 places a strong emphasis on target (reflected in 2D as well-defined and separated clusters). The default of 0.5 balances the weighting equally between data and target, see Fig S4. Essentially umap takes the labels as a separate metric space (with a categorical distance on it) and tries to fold the two data and labels together by performing an intersection of the simplicial sets. The parameter target_weight provides some level of balance between how much weight is applied to the label vs data. A target_weight of 1.0 will put almost all the weight on the labels, while a target_weight of 0.0 will weight as much as can be managed in favour of the data.

#### n_neighbors

For supervised UMAP embeddings it is recommended to use a higher value of *n_neighbors*. However, if the user is only interested mostly in very local information can use low values of n_neighbors.

#### target_metric

By default, the *target_metric* is set to ‘categorical’, we do not recommend changing this parameter to another metric such as euclidean.

For more UMAP parameters to tune the embeddings please refer to the work of McInnes et al., 2018.

The important parameters for mb-PHENIX function are:

#### knn and decay

For the adaptive kernel to construct the Markovian matrix, the user chooses a *knn* value that is the number of nearest neighbors from which to compute kernel bandwidth. The parameter decay is the decay rate of kernel tails. By default, the decaying kernel is set to 1, we recommend default setting for decay. For small datasets we recommend a set *knn* value sufficient to avoid over-smoothing to other clusters but not too small to alter the connectivity of data as a graph.

#### t

For the diffusion process, the parameter *t* (diffusion time) is the power value to which the Markovian matrix is powered. This sets the level of diffusion.

The *knn* and *t* values need to be sufficient to build a complete graph (considering the class) and less to avoid over-smooth taxa to other distinct classes.

In Fig S3 there is a code example of how to apply this in Google colab for an easy implementation.

### S4 Imputation of microbiota data and the analysis of the imputed data

Here we show two examples of imputation methods mb-PHENIX can be used for 16S RNA and WGS datasets and some other methods of imputation of high-dimensional data. We compared mb-PHENIX, MAGIC and mbImpute.

In a Mexican population with prediabetes and diabetes, Diener et al 2019 studied reported in 16S amplicon sequence data from a cohort of 405 participants were divided into 6 groups: healthy individuals, IFG, IGT, IFG+IGT, T2D. With 16s rRNA metagenomic data from each participant (n=405). In this investigation, the authors reported that *Escherichia* and *Veillonella* were associated with T2D progression, along with biochemical measures of blood glucose and insulin-related measures, respectively. At the same time, *Blautia* and the *Anaerostipes* decreased with T2D development. Non-imputed data follow an unimodal distribution per taxon, all towards the 0 value, due to the zero inflated nature of the microbiota data (Fig S11).

After a supervised imputation with mb-PHENIX using sUMAP embedding, In Fig S6, we find unique multimodal distributions per distinct taxon, with different transition states of TD2 clusters represented by different peaks, consistent with the known taxon in these microbiome subsets, all reported in Diener et al 2019. Interesting *Akkermansia* was not reported in that work as to be involved in T2D or in its intermediate’s states of this disease. Our approach indicates that there is a strong signal in Fig S6 that probably a specific strain of *Akkermansia* is correlated in T2D and IFG.

The COVID 19 microbiota data set was taken from Zho et al, 2020. It consists of 15 patients with COVID-19 (cases), 15 healthy control individuals and 6 patients with non-covid-19 related pneumonia (control). The cases’ fecal samples were taken during hospitalization at different times. Then, they performed WGS to obtain taxonomic profiles for each sample. We downloaded the data from NCBI bioproject PRJNA624223. With these sequences we extracted the information to perform three different imputation methods, MAGIC, mbimpute and mb-PHENIX. In the figure S5, we show the most variable taxa, only mb-PHENIX recovers more taxa involved in the different stages of the COVID-19 disease. For example, with our approach (Fig S11), we found that *Bordetella, fusobacterium* and *Slackia* as putative novel markers that are involved in the critical stage of the COVID-19 (work in development) disease and systemic infection.

## Supplementary section 1

**Fig S1.**
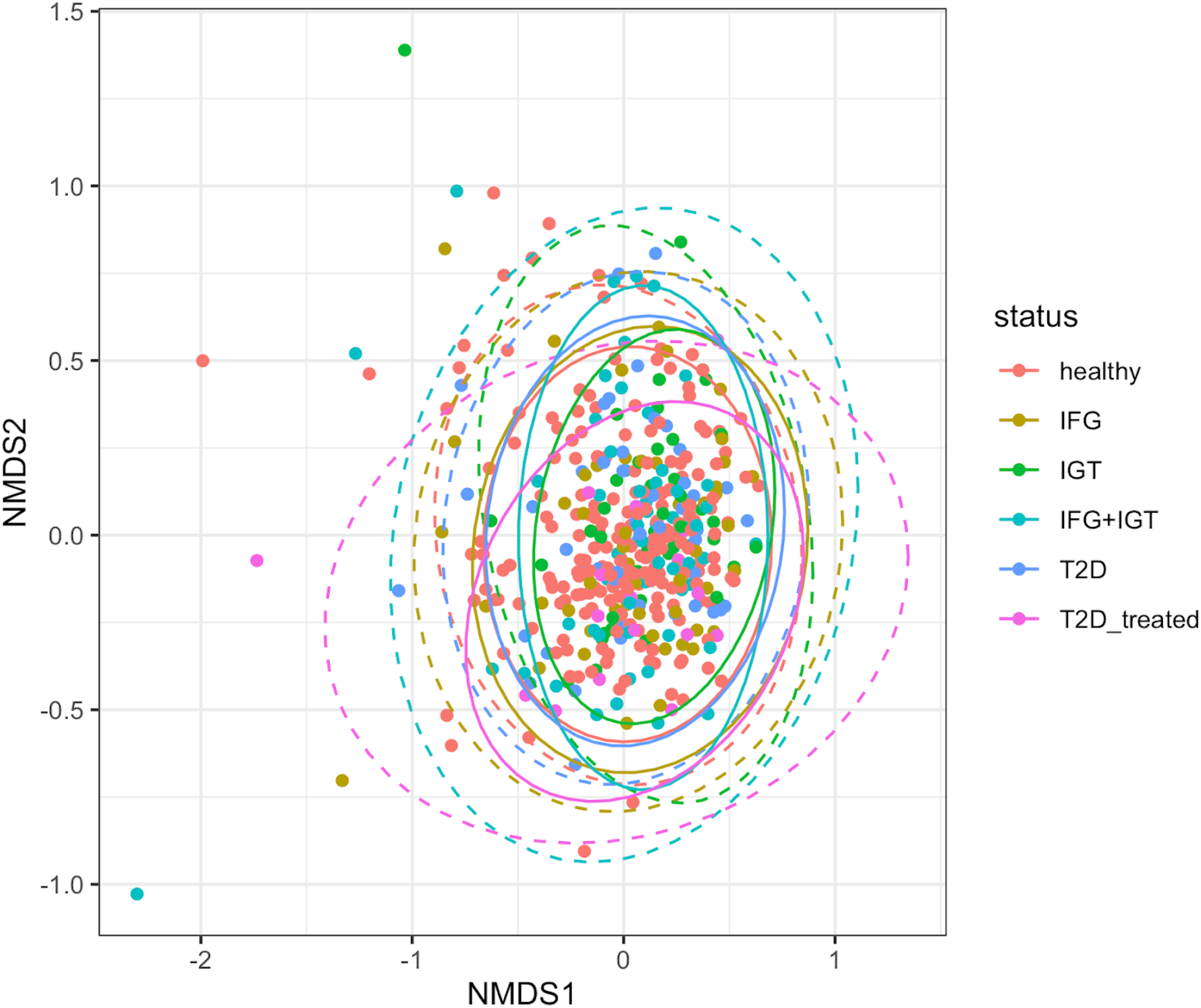
NMDS plot of T2D Mexican cohort. Progressive Shifts in the Gut Microbiome Reflect Prediabetes and Diabetes Development in a Treatment-Naive Mexican Cohort. We can observe that there is not a well-defined cluster structure among the Healthy, IFG, IGT, IFG+IGT, T2D and T2D treated groups. This is because of the high-heterogeneity of microbiota and the detrimental effects of the technical noise and high-dimensionality. The groups are: Healthy, IFG(impaired glucose tolerance), IGT(impaired fasting glucose), IFG+IGT(impaired glucose tolerance and impaired fasting glucose), T2D(diabetes type 2) and T2D treated groups. NMDS= Non-metric Multi-Dimensional Scaling.

**Figure S2:**
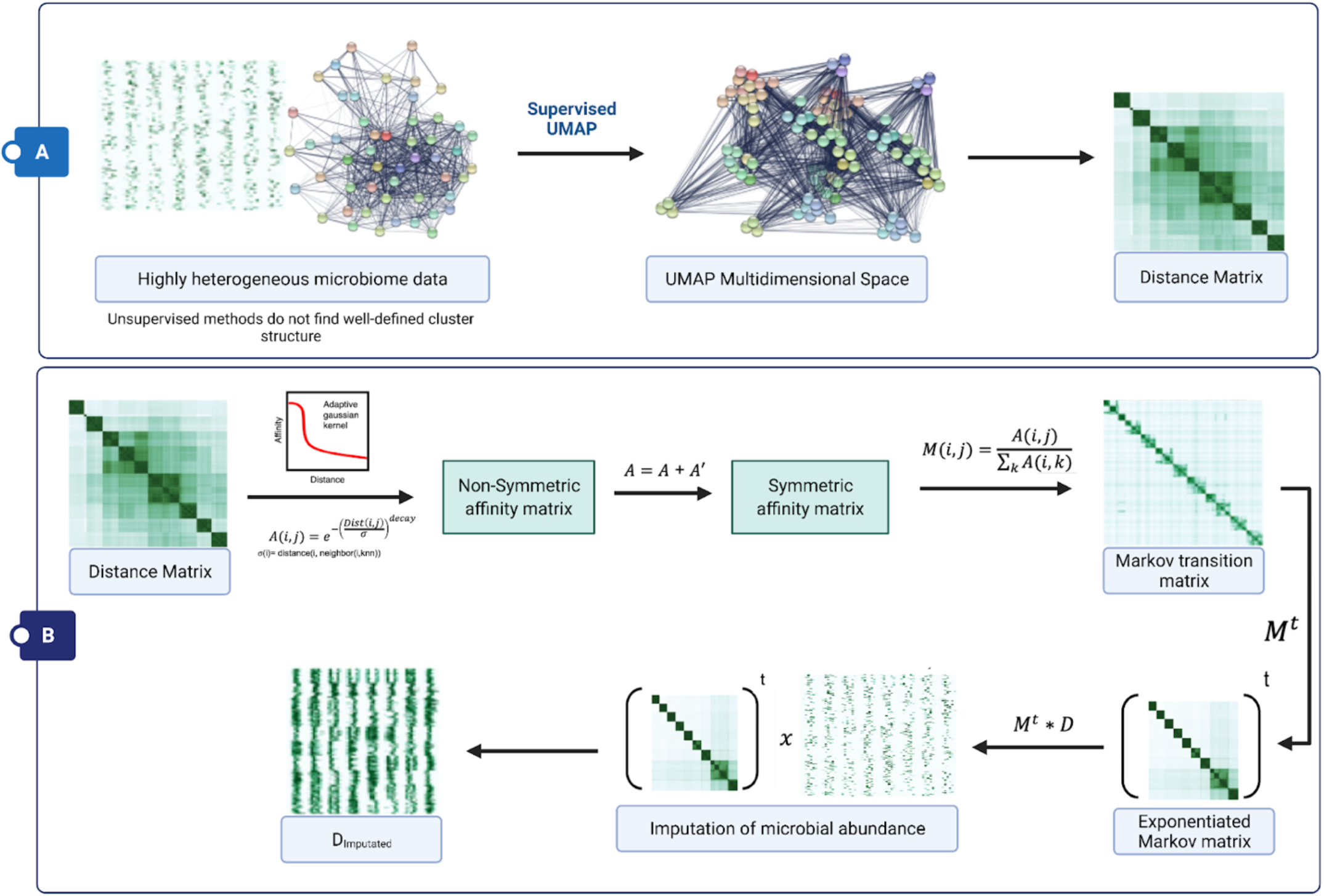
The imputation process using mb-PHENIX. The mb-PHENIX imputation approach for microbiota data consists of two main parts. A) The construction of the distant matrix: The distance matrix is computed from the UMAP space. The method mb-PHENIX is characterized by applying UMAP in a supervised regiment. B) The diffusion maps for imputation: For the imputation process using diffusion maps consist of several steps: 1. Construction of Markov transition matrix from the distant matrix; mb-PHENIX uses the adaptive kernel to generate a non-symmetric affinity matrix, it is symmetrized and then is normalized to generate the Markov transition matrix. 2. Diffusion process; the Markov transition matrix *M* is exponentiated to a chosen power t (random walk of length t named “diffusion time”). 3. Imputation; imputation of gene expression consists in multiplying the exponentiated Markov matrix (*M*^*t*^) by the taxa count data to obtain an imputed and denoised taxa count matrix. The Equations are described in more detail in (Padron-Manrique et al., 2022).

**Fig S3.**
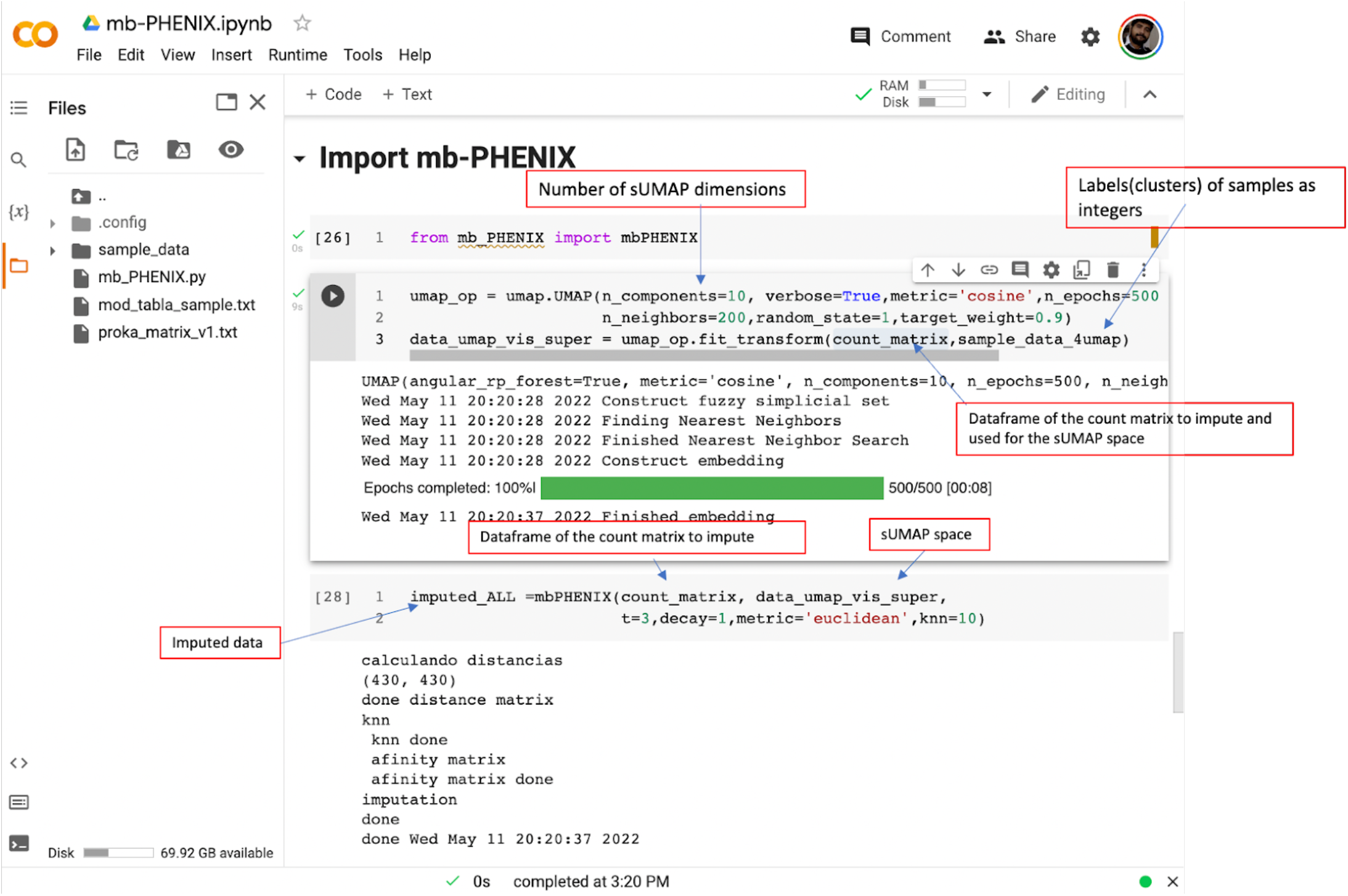
Code implementation of MB-PHENIX. The input of mb-PHENIX is an abundance matrix where the taxa (or ASV/OTUs ID) are columns and rows are samples. This input matrix needs to be a PANDAS dataframe.

**Fig S4.**
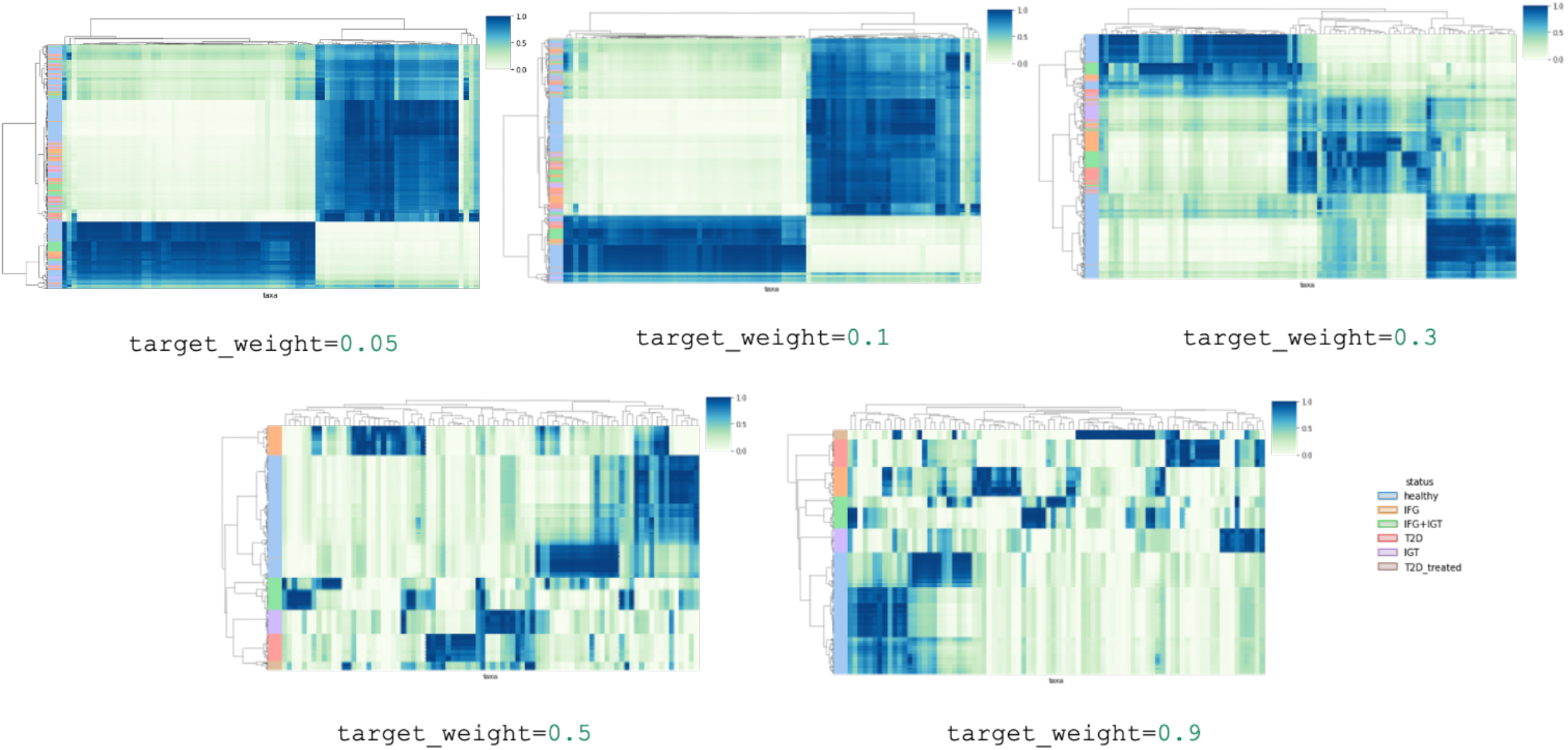
Heat map of the highly variable taxa imputed by mb-PHENIX using different values of the parameter *target_weight*. The data of the Mexican cohort (Diener et al., 2019). We observed increasing the values of target_weight with mb-PHENIX the imputed microbiome data recovered cluster structure of the different stages of diabetes.

**Fig S5.**
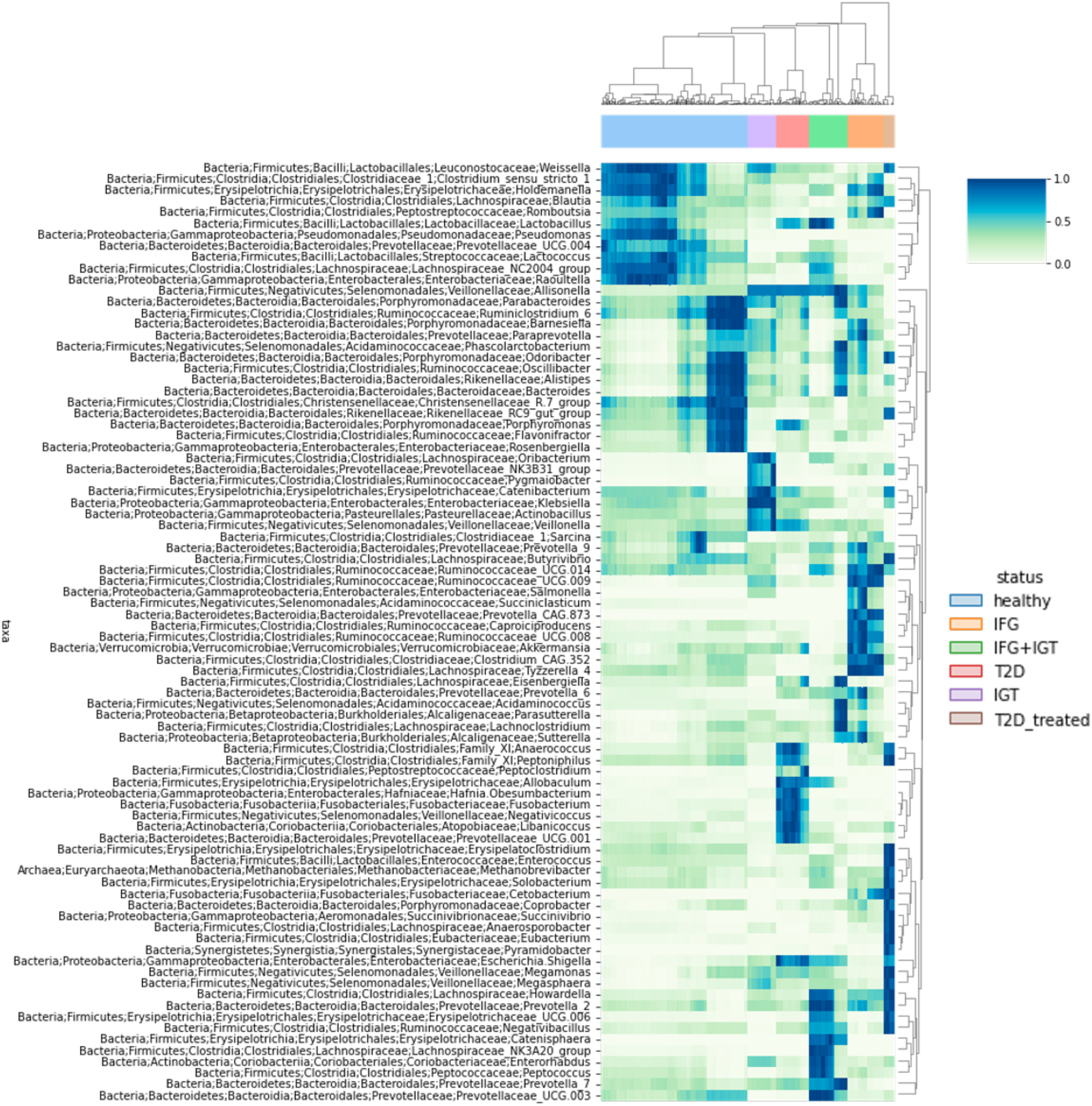
The method mb-PHENIX reveals multinomial distribution of taxon in the progregression of T2D microbiota data. Plot of the imputed taxa matrix as a hierarchically-clustered heatmap. We visualize the most variable taxa(95 percentile) data after supervised mb-PHENIX imputation.

**Fig S6.**
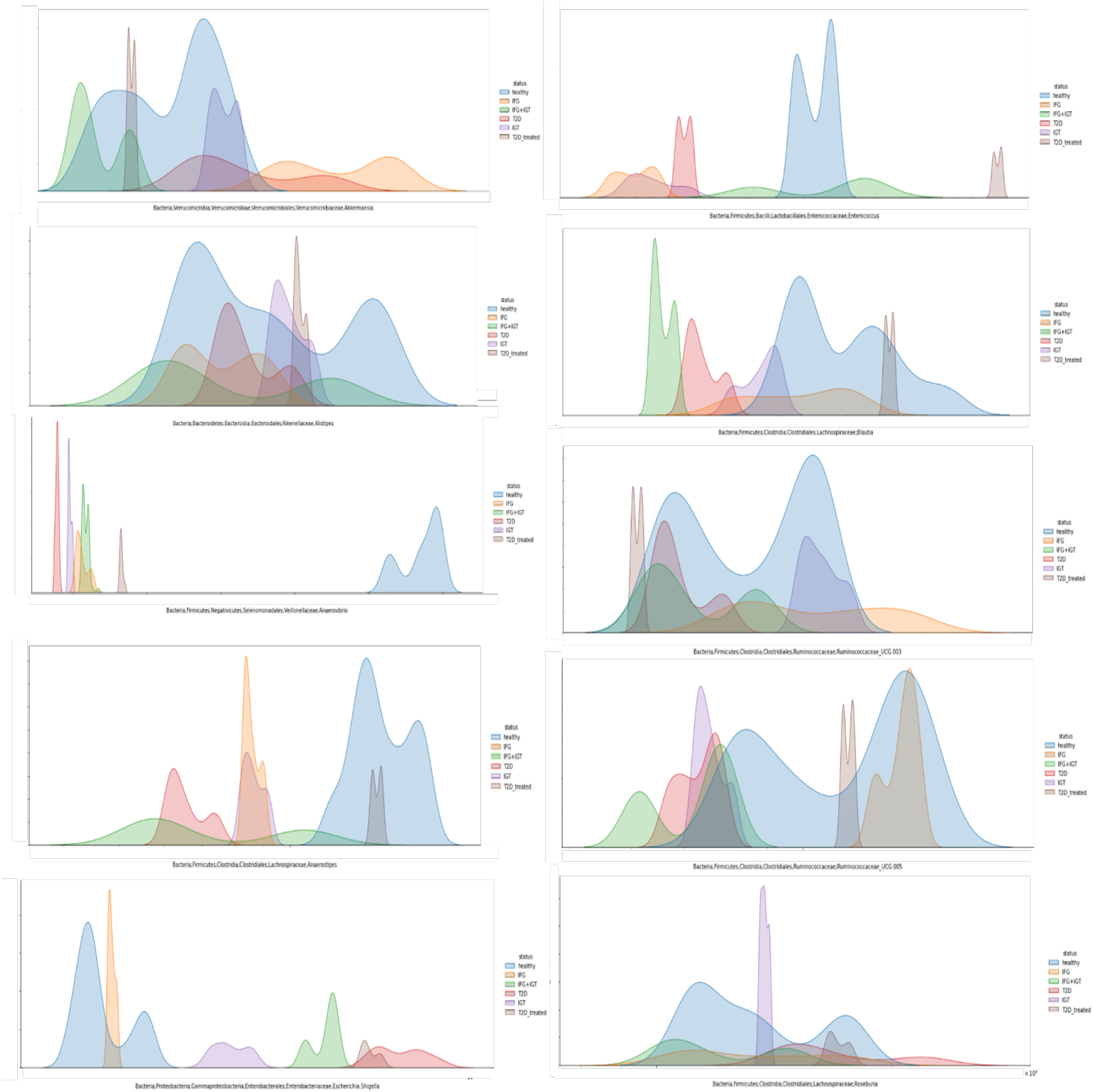
The method mb-PHENIX reveals multinomial distribution of taxon in the progression of T2D microbiota data. Histograms by state clusters for taxa (*Akkermansia, Enterococcus, Alistipes, Blautia, Anaerostipes, Anaerovibrio, Ruminococcaceae_UCG*.*00# Escherichia shigella and Roseburia*) calculated using Kernel Density Estimation (KDE) on the data after supervised mb-PHENIX imputation. After supervised mb-PHENIX, we observed unique multimodal distributions per taxa, with different transition states of TD2 clusters represented by different peaks, consistent with the known taxa in these microbiome subsets. TD2= Type 2 Diabetes, IFG= Impaired Fasting Glucose, IGT = Impaired Glucose Tolerance, T2D_treated = Treated Diabetes, IFG+ IGT = Impaired Fasting Glucose and Impaired Glucose Tolerance.

**Fig S7.**
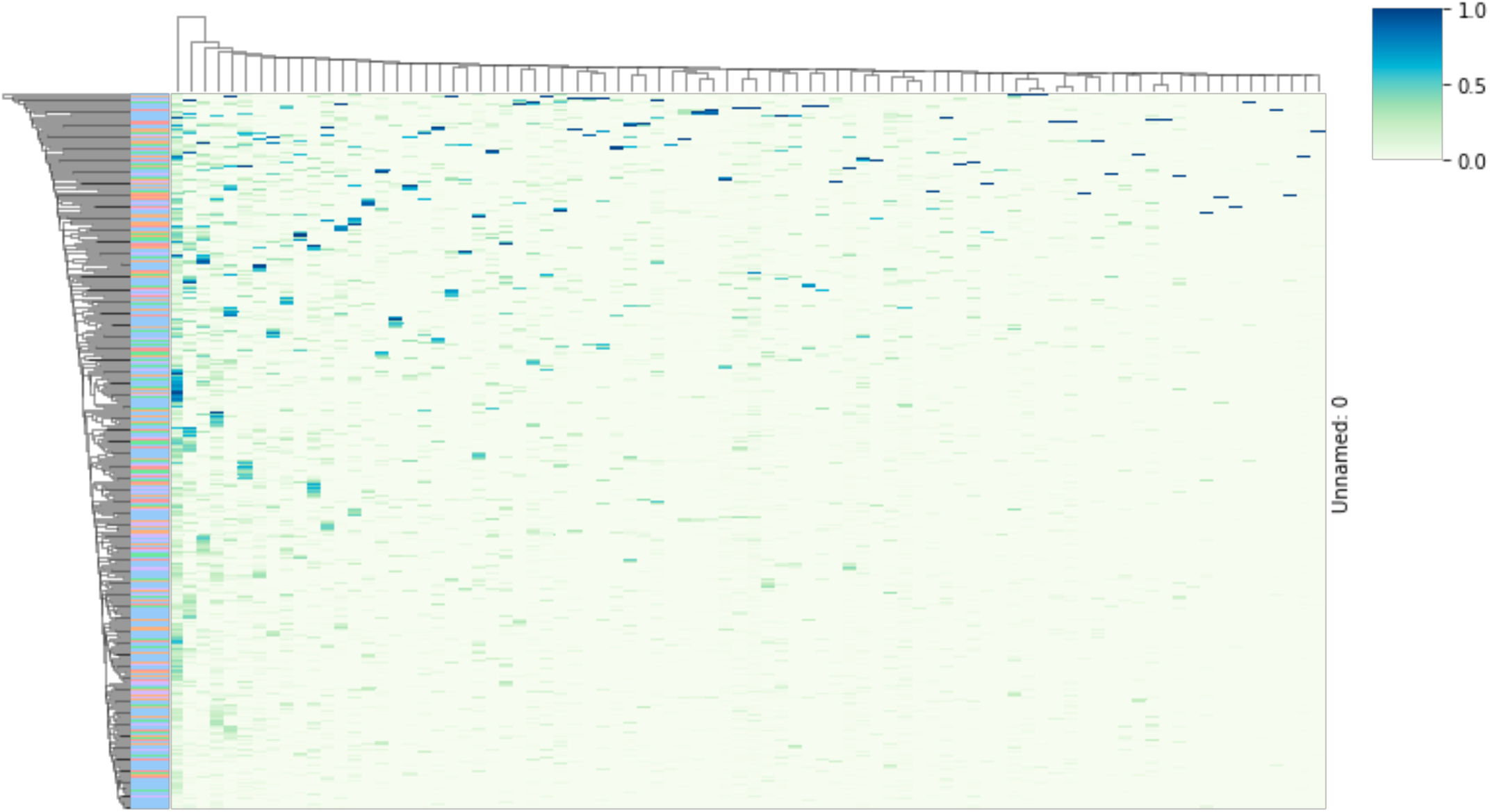
Hierarchically-clustered analysis of the Imputed data with mbImpute. Plot of the taxa matrix imputed with mbImpute as a hierarchically-clustered heatmap. We visualize the most variable taxon(95 percentile). There are not well-defined clusters using mbImpute.

**Fig S8.**
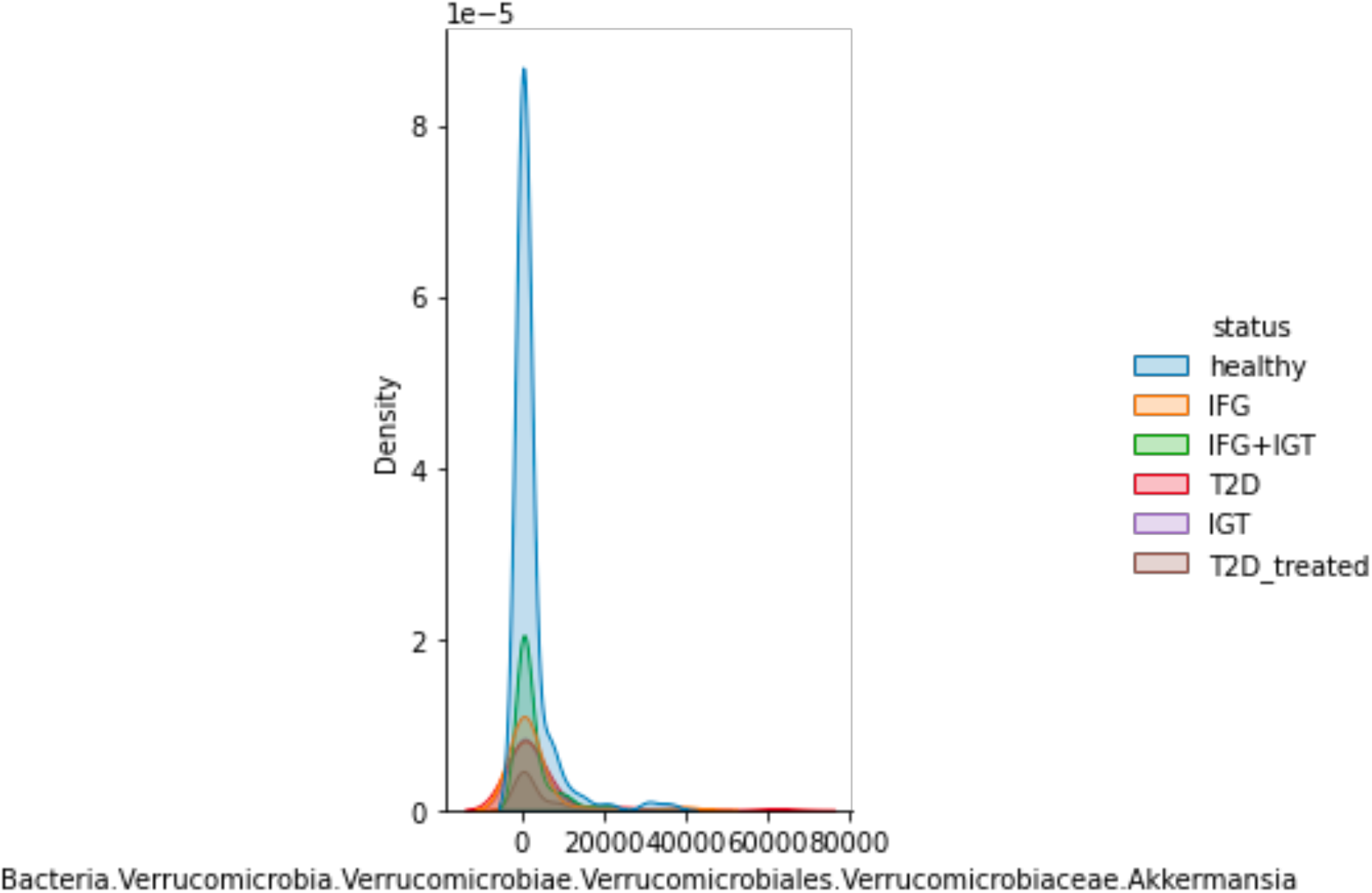
Histogram of the imputed data with mbImpute. Histograms by state clusters for *Akkermansia* calculated using Kernel Density Estimation (KDE) on the data after unsupervised MAGIC imputation. After unsupervised mbImpute, we observed an uni multimodal distribution of the *akkermansia* taxon, with different transition states of TD2 clusters represented by different peaks. There are no clear differences among different stages of TD2 on the imputed data with MAGI. TD2= Type 2 Diabetes, IFG= Impaired Fasting Glucose, IGT = Impaired Glucose Tolerance, T2D_treated = Treated Diabetes, IFG+ IGT = Impaired Fasting Glucose and Impaired Glucose Tolerance.

**Fig S9.**
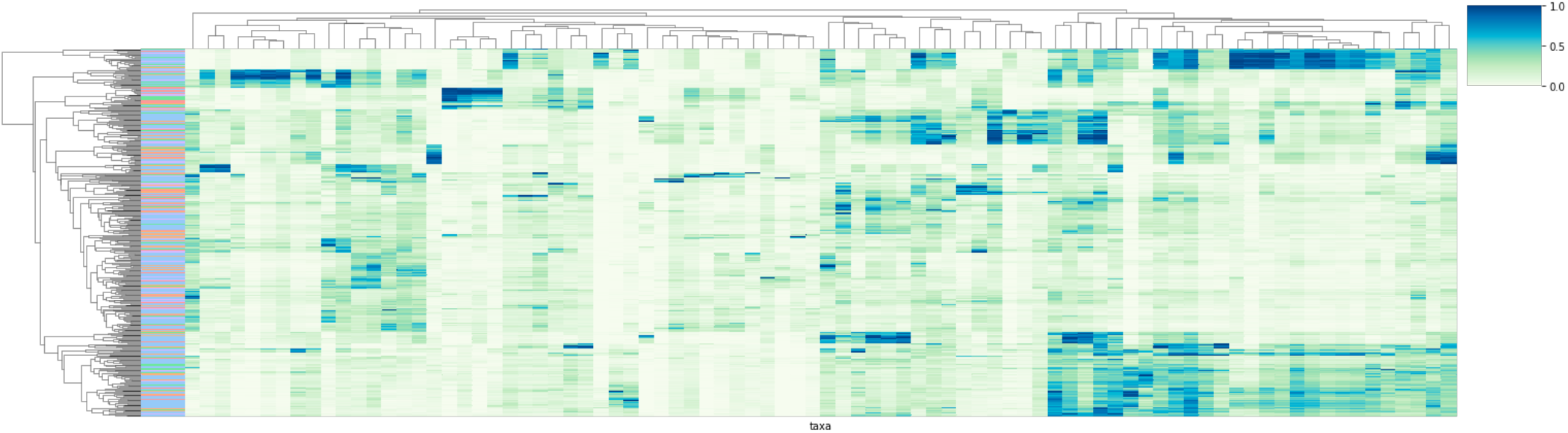
Hierarchically-clustered analysis of the Imputed data with MAGIC. Plot of the taxa matrix imputed with mbImpute as a hierarchically-clustered heatmap. We visualize the most variable taxon(95 percentile). There are not well-defined clusters using MAGIC.

**Fig S10.**
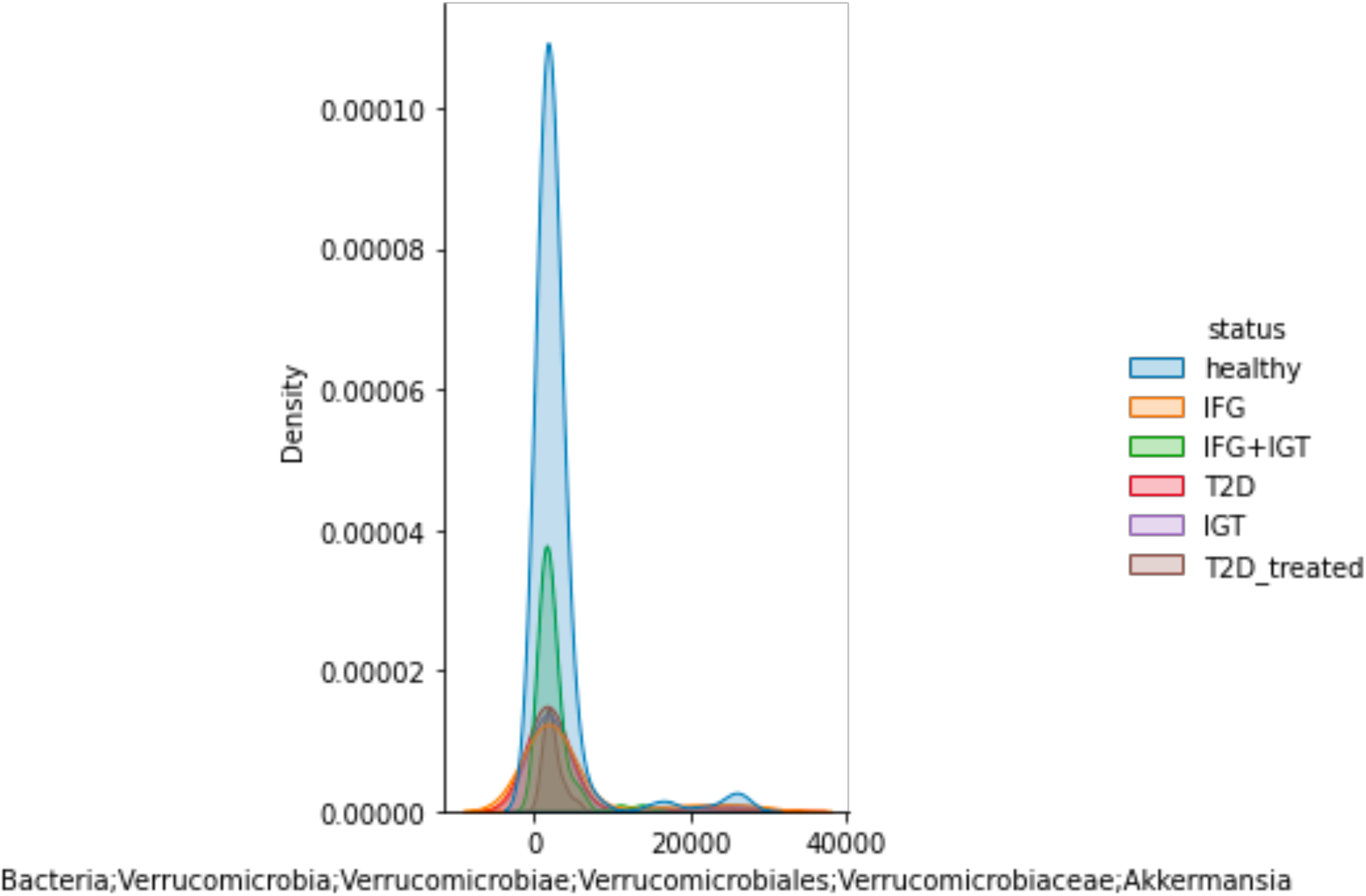
Histogram of the imputed data with MAGIC. Histograms by state clusters for *Akkermansia* calculated using Kernel Density Estimation (KDE) on the data after unsupervised MAGIC imputation. After unsupervised MAGIC imputation, we observed an uni multimodal distribution of the akkermansia taxon, with different transition states of TD2 clusters represented by different peaks. There are no clear differences among different stages of TD2 on the imputed data with MAGI. TD2= Type 2 Diabetes, IFG= Impaired Fasting Glucose, IGT = Impaired Glucose Tolerance, T2D_treated = Treated Diabetes, IFG+ IGT = Impaired Fasting Glucose and Impaired Glucose Tolerance.

**Fig S11.**
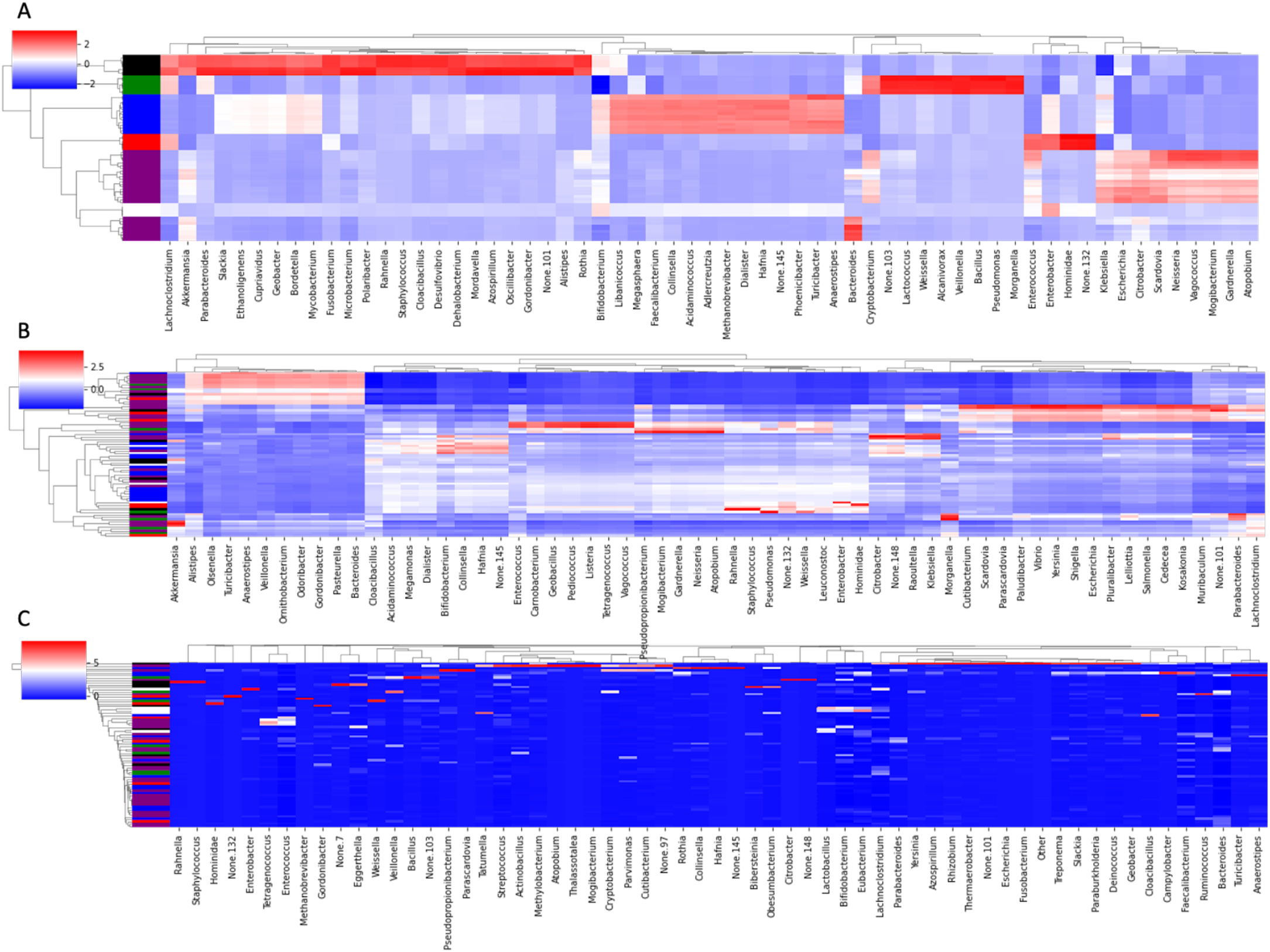
Heat map of the imputed highly variable taxa using the COVID-19 data. We imputed it with MAGIC, mbImpute and mb-PHENIX. Cluster colors: Black= severity level 4(critical condition), green= 3, purple = severity level 2, white= severity level 1, Blue= Non Covid(healthy) and red= neumonia. Imputation using A) mb-PHENIX, B) MAGIC and C) mbImpute

**Fig S11.**
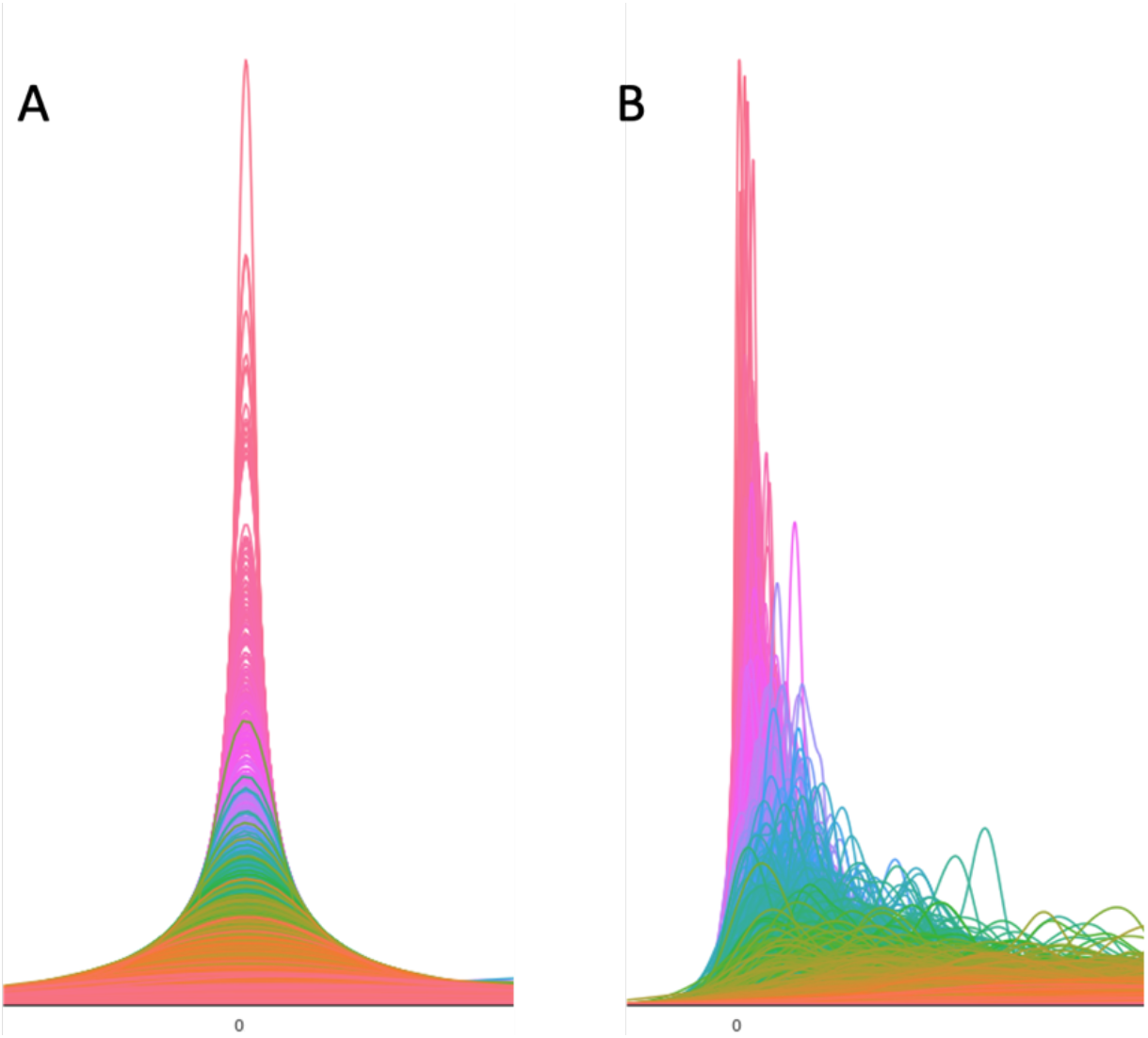
Distribution of the taxon counts. The mb-PHENIX method reveals multimodal taxon distribution.. Histograms of taxa data from all samples computed using kernel density estimation on the data before (A) and after (B) mb-PHENIX imputation. Due to the high amount of missing taxa, most density is concentrated unimodally at zero. Before mb-PHENIX we observed multimodal distributions per ASV.

